# Multi-omic integration using interpretable machine learning reveals genetic mechanisms of trait variation and phenotypic plasticity in switchgrass

**DOI:** 10.64898/2026.03.06.710154

**Authors:** Paulo Izquierdo, Xiaoyu Weng, Thomas E. Juenger, Jason Bonnette, Yuko Yoshinaga, Chris Daum, Anna Lipzen, Kerrie Barry, Matthew Blow, Melissa D. Lehti-Shiu, David B. Lowry, Shin-Han Shiu

## Abstract

Uncovering the genetic architecture of quantitative traits is challenging because polygenic control yields small individual gene effects and because gene–gene and genotype-by-environment interactions add further complexity. To understand the genetic basis of polygenic traits and their plasticity across environments, we integrated genome-wide SNPs and RNA-seq transcript data with interpretable statistical and machine learning models in a switchgrass (*Panicum virgatum*) diversity panel grown at contrasting field sites in Michigan and Texas. Notably, in addition to single environments, our trait prediction models were able to predict phenotypic differences, across environments i.e., plasticity. By interpreting trait prediction models with explainable machine learning methods, we identified genes that are the most predictive of flowering time and annual biomass production across environments, based on their associated gene expression levels and nearby SNPs. This approach recovered canonical flowering regulators and revealed novel, environment-specific candidate flowering genes. Further, transcriptome models consistently recovered more switchgrass genes homologous to experimentally validated genes in Arabidopsis and rice than SNP-based models. Feature interaction scores from the models also allow the identification of trait- and environment-dependent gene–gene interactions, where flowering time showed stronger and more abundant interactions than biomass. While some of the interactions identified are consistent with the link between flowering time and yield, most are novel predictors that need to be further evaluated. These findings provide insight into the genetic basis of trait variation and plasticity across environments and help prioritize candidate genes for functional validation and cultivar improvement.

## Introduction

Polygenic traits are controlled by multiple loci with varying effects. The contributions of these loci are further modulated by interactions with other genes (i.e., gene–gene [G×G] or epistatic interactions) and the environment (i.e., genotype-by-environment [G×E] interactions), complicating genetic mapping and the identification of loci underlying traits. Understanding the genetic mechanisms of G×E is important because it represents a key source of genetic variation that can be leveraged for climate-resilient breeding. For example, G×E can reveal conditionally neutral loci that confer trait advantages in specific environments without incurring trade-offs in others^1^, offering a pathway to harness adaptive plasticity in crops. However, despite a large body of research on trait architecture, most studies have been conducted in single environments. As a result, both the contributions of G×E interactions to trait variation and their underlying genetic mechanisms remain largely unresolved^2,3^.

To dissect the genetic mechanisms that govern trait variation within environments and the plasticity arising from G×E, it is advantageous to study species spanning broad environmental gradients^4^. One such species is switchgrass (*Panicum virgatum*), a native North American perennial grass with a broad latitudinal range and pronounced upland, lowland, and coastal ecotypic differentiation^5,6^. This distribution has enabled the establishment of a common-garden diversity panel across a latitudinal gradient, which has revealed climate adaptation among genotypes^6^. Despite the major phenotypic differences between ecotypes, earlier findings revealed that allelic variation within ecotypes could either increase or decrease traits such as biomass, depending on the environment^7^.

In addition to the availability of genetic resources facilitating studies of G×E, recent advances in high-throughput genotyping, transcriptomics, and data-driven modeling are opening new opportunities to connect predictions with underlying biological mechanisms across multiple environments^8,9^. Integrating omics data with interpretable machine learning, or explainable artificial intelligence^10^, has recovered benchmark genes and revealed novel candidates for flowering-time control in maize (*Zea mays*) and *Arabidopsis thaliana*^11,12^. Moreover, it is possible to estimate interactions between predictors using approaches such as SHAP (SHapley Additive exPlanations)^13^. SHAP quantifies the contribution of each feature to individual predictions in a consistent and additive manner, enabling the interpretation of complex, non-linear models. These analyses have been used to generate model-based hypotheses of epistasis in both humans and *Arabidopsis*, some of which have been supported by experimental evidence^12,14^. Together, these approaches provide a framework for generating mechanistic hypotheses and facilitating biological discovery across environments.

Despite these advances, we still lack a clear understanding of how genomic and transcriptomic signals contribute differently to complex trait variation and plasticity across environments. It remains unresolved whether transcriptome-based models capture distinct, environmentally responsive information compared to genotype-based models. Moreover, we do not yet know how gene effects are modulated by genetic background and environment, or how gene–gene interactions change across conditions. Addressing these gaps is critical for translating predictive models into biological hypotheses about trait regulation and plasticity.

To address these gaps, we leveraged phenotypic data from a switchgrass diversity panel grown in common gardens in Texas (TX) and Michigan (MI) to dissect the genetic basis of flowering time and biomass in two contrasting environments. We generated machine learning models to predict trait variation within each site and plasticity between sites using single nucleotide polymorphisms (SNPs) identified from existing whole-genome sequencing data^6^ and newly generated transcriptomic variation data. To interpret the models, we identified genes and gene-gene interactions important for trait predictions and compared them to genes homologous to experimentally validated genes associated with flowering and biomass in Arabidopsis and rice. Finally, we evaluated how gene effects are influenced by genetic background and environmental context, and how genes contribute to trait plasticity across environments.

## Results and discussion

### Environmental variation drives phenotypic and transcriptional plasticity in switchgrass

To investigate the extent to which traits are influenced by environmental variation, we measured six traits (green-up, emergence, flowering time, tiller count, panicle height, biomass) for 426 tetraploid genotypes from a switchgrass diversity panel grown in Texas (TX) and Michigan (MI), which differ markedly in latitude (**see Methods**). Phenotypic data were collected at both locations during the 2021 growing season. To quantify phenotypic plasticity across environments, we calculated Pearson correlation coefficients (PCCs) between trait values across environments. PCC values near ±1 indicate low plasticity (i.e., consistent trait rankings across environments and little G×E), and values near 0 indicate high plasticity (i.e., strong G×E effects)^15^. PCCs ranged from –0.13 (*p*<8.3×10^-3^) for green-up to 0.75 (*p*<2.2×10^-16^) for flowering time, indicating variable plasticity across traits (**Supplementary Fig. S1**). Next, we evaluated the extent to which phenotypic variance across environments could be explained by G, E, and G×E. We found that early developmental traits were most strongly influenced by E (green-up = 98%, emergence time = 73%), followed by mid-stage traits (flowering time = 68%), and, finally, mature traits (plant height = 16%, tiller count = 14%, biomass = 26%). In contrast, the contributions of G×E were higher for mid-stage and mature traits **(Fig. 1a)**. This pattern suggests that environmental effects contribute most to phenotypic variation in early developmental stages, whereas genetic factors contribute more as plants mature. It should be noted that, because only a single replicate of each genotype was established at each site, fine-scale environmental variation within field sites may have contributed to a portion of the phenotypic variation attributed to G×E.

**Figure 1.**
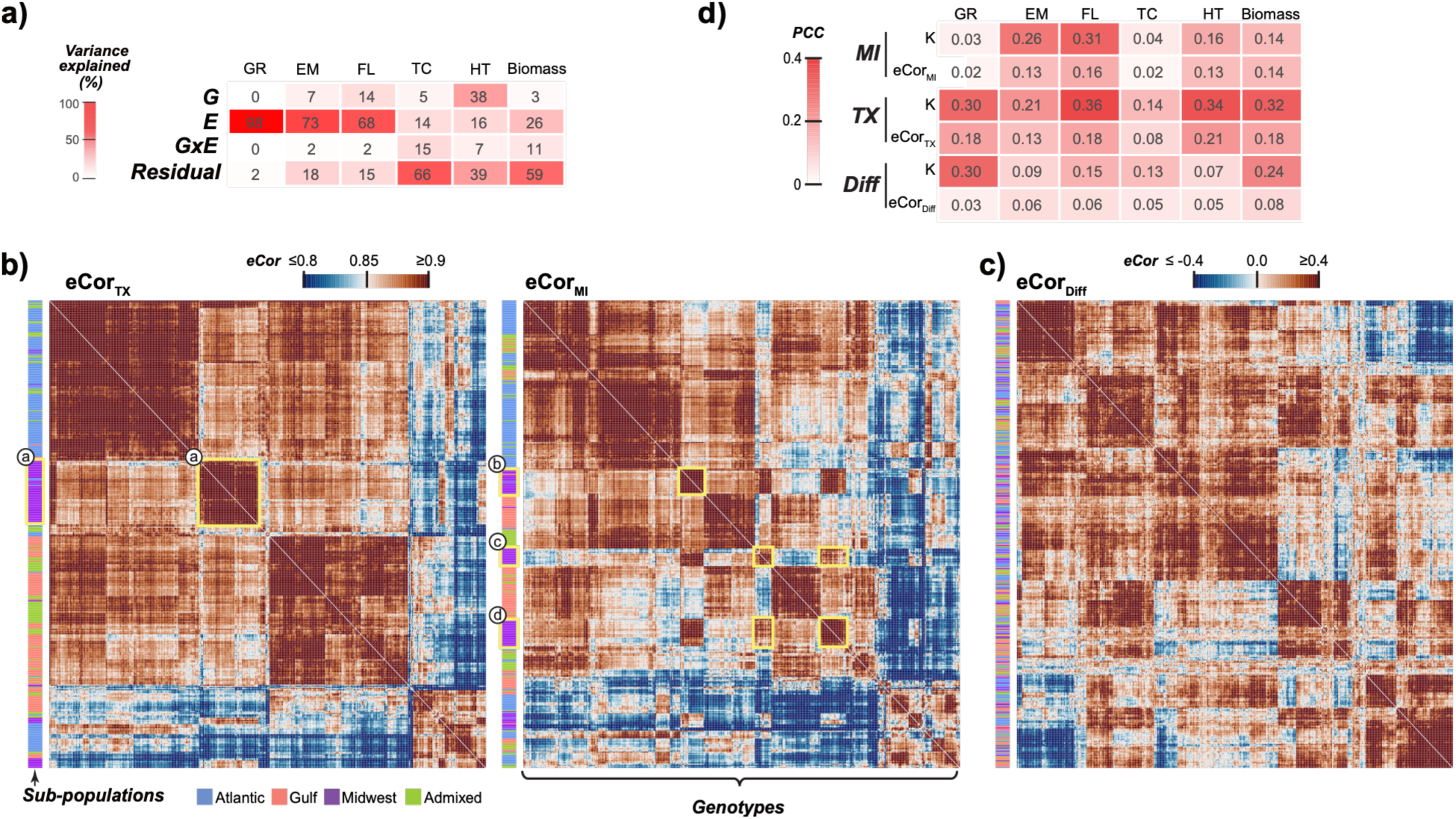
Relationships among phenotypes, gene expression, and genomic similarity across environments. **(a)** Proportion of phenotypic variance explained by genotype (G), environment (E), and genotype-by-environment interaction (G×E) for six traits: green-up (GR), emergence (EM), flowering time (FLT), tiller count (TC), panicle height (HT), and log-transformed biomass. Pairwise expression correlation (eCor) among all genotypes based on transcriptomic data from **(b)** TX and MI (yellow rectangles highlight subpopulation-specific genotype clusters that shift across environments) and **(c)** gene expression differences between TX and MI (Diff), with heatmap colors ranging from blue (low) to white (intermediate) to brown (high) indicating the degree of correlation. **(d)** Pearson correlation coefficients (PCCs) between genomic relationship (K) or eCor based on TX, MI, or Diff transcript levels and phenotypic values in TX or MI or the difference in values between TX and MI (Diff). Heatmap colors ranging from white (zero) to red (positive) indicate PCC. Note that K calculated was using all genotypes, and the same value was used for all PCC calculations.

Given the environmental and G×E effects on trait variation, we next asked whether gene expression profiles similarly reflect environment-specific responses across genotypes. To test this, we collected transcriptome data from each genotype at both field sites (**see Methods**). Transcriptomic similarity among genotypes was measured using expression correlation (eCor), calculated as the PCC of gene expression levels. Clustering patterns of eCor differed between environments, indicating that while genotypes from similar genetic backgrounds (i.e., subpopulations; **Fig. 1b**) tend to cluster together, they also exhibit environment-specific transcriptional responses (e.g., clusters a–d; **Fig. 1b**). To investigate this further, we asked whether expression differences across environments (DiffExp = TX − MI) reflect genetic similarity. We calculated DiffExp as the difference in log₁₀(TPM + 1)-transformed expression values between Texas and Michigan, which can be interpreted as a measure of relative expression change between environments. If gene expression responses are genotype-driven, we would expect genetically similar individuals to show similar expression changes and cluster by subpopulation. Conversely, if responses are environment-specific, expression differences should not follow subpopulation structure. We observed no clear clustering by subpopulation in expression plasticity (**Fig. 1c**), indicating that across-environment transcriptional responses are primarily driven by environment rather than ancestry.

Having observed that gene expression within environments tends to cluster genotypes by genetic background, but transcriptional responses across environments do not, we next asked whether genetic similarity correlates with transcriptomic similarity. We found that genomic relatedness (kinship) derived from SNP data (**see Methods**) was significantly correlated with eCor within both TX (r = 0.38, *p*<2.2×10⁻¹⁶) and MI (r = 0.37, *p*< 2.2×10⁻¹⁶), but was not correlated with Diff_exp_ (r = 0.07), consistent with clustering patterns observed in **Fig. 1b–c**. These results reinforce the observation that transcriptional responses across environments are decoupled from genetic similarity and instead largely shaped by environment. We assessed the relationships among kinship, eCor, and phenotypic similarity within each environment to determine the extent to which genetic relatedness is associated with similarity in expression and phenotypic profiles. Overall, both kinship and eCor showed stronger correlations with trait values in TX than in MI (**Fig. 1d**), consistent with the higher proportion of trait variation explained by genetic effects (heritability) in TX for all traits except tiller count (**see next section**). Next, we tested whether variation in trait plasticity (Diff_trait_ = TX – MI) could be explained by either kinship or Diff_exp_. Kinship partially explained variation in Diff_trait_ for some traits, notably biomass (r = 0.24) and green-up (r = 0.30), while Diff_exp_ showed low correlation with trait plasticity across all traits (r ≤ 0.08; **Fig. 1d**). These results demonstrate that genetic variants and gene expression contribute differently to trait variation and plasticity, reflecting distinct biological and environmental influences.

### Omics data enable the prediction of traits in single environments and trait plasticity

To evaluate the extent to which transcriptomic and genetic variant (i.e., SNP) data predict switchgrass traits across environments, we built genomic prediction models using transcriptomic (T), SNPs, or both (S+T) as input features (**Fig. 2**). SNPs remain constant for each genotype across environments, whereas transcriptomic data capture environmentally responsive changes in gene expression. Because trait variation involves both linear and nonlinear relationships among genes and traits^16^, we compared a linear (Bayesian Ridge Regression, BRR) with a nonlinear model (Extreme Gradient Boosting, XGBoost).

**Figure 2.**
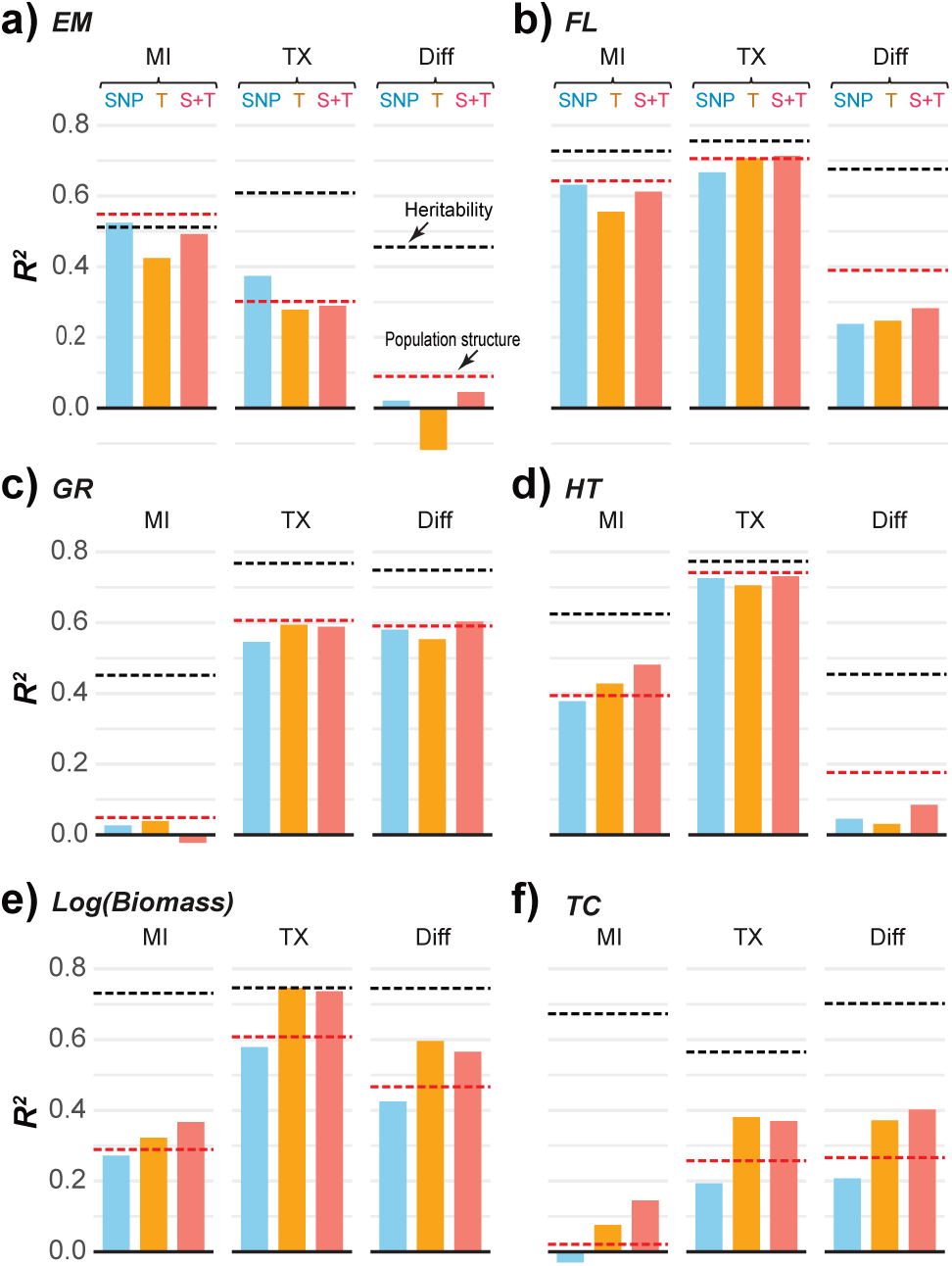
Model performance across locations and traits. **(a-f)** XGBoost model performance (measured as *R²* using a testing set that has not been subjected to the training process) for six traits using SNPs, transcriptomic (T), and their combination (S+T). Model performance based solely on population structure (first five principal components based on SNPs) is indicated by red dotted lines. Heritability estimates (black dotted lines, 0–1 scale) are shown for reference and are not equivalent to R².

Model performance varied by trait but was consistent across algorithms and data types (**Supplementary table S1**). XGBoost models, except for biomass and tiller count models, did not outperform predictions based on the first five principal components of SNP data, which were used to approximate population structure (**Fig. 2**). Although we found that kinship correlated more strongly with phenotypic variation than eCor **(Fig. 1d)**, both transcriptomic- and SNP-based models achieved comparable predictive performance. As expected, traits with higher correlations with genetic similarity (i.e., kinship or eCor; **Fig. 1d**) were predicted with higher accuracy (e.g., flowering time and biomass; **Fig. 2**). Notably, transcriptomic-based models of biomass outperformed both SNP- and population structure-based models across sites, approaching the accuracy of heritability estimates in TX (**Fig. 2e**).

Because E and G×E effects together accounted for 20–98% of phenotypic variance across environments (**Fig. 1a**), we hypothesized that transcriptomic plasticity could serve as a predictor of trait plasticity (i.e., differences in trait values across environments). To test this, we trained models using SNP data, which remain constant across environments, and transcriptomic data, where plasticity was estimated as the difference in expression between TX and MI. We found that Diff_exp_ models were more predictive than SNP models for trait plasticity in biomass (R² = 0.60 vs. 0.42) and tiller count (0.37 vs. 0.21), showed comparable performance for flowering time (0.25 vs. 0.24), and were slightly less predictive for green-up (0.55 vs. 0.58). For emergence and panicle height, however, both data types yielded low predictive performance (R² < 0.1), suggesting that the available features were insufficient to capture the plasticity of these traits. We further hypothesized that trait plasticity would be more predictable for traits with low phenotypic correlation between environments. Consistent with this hypothesis, difference-based SNP and transcriptomic models showed predictive performance comparable to or greater than at least one single-environment model for traits with low cross-environment correlation, such as green-up (**Fig. 2c**, r = –0.13, p < 8.3 × 10⁻³) and tiller count (**Fig. 2f**, r = –0.08, p = 0.1) (**Supplementary Fig. S1**). In contrast, for traits with high cross-environment correlation, such as flowering time and panicle height (**Fig. 2b,d**, r > 0.68, both p < 2.2 × 10⁻¹⁶), difference-based models generally exhibited lower predictive performance than single-environment models. Together, these results indicate that SNP- and transcriptomic-based models differed in their predictive performance across traits and environments, with each data type showing advantages in specific contexts. These differences suggest that SNPs and gene expression features capture partially distinct sources of biological information underlying trait prediction.

### Transcript features enhance recovery of known and novel genes underlying trait variation and plasticity

Because SNP- and transcriptomic-based models yielded comparable predictive accuracy, we tested whether integrating both data types would improve performance. Multi-omic models that included both transcriptomic and SNP features outperformed either model alone in only 15% of cases, with modest gains (R² ≤ 0.07); the largest improvement was observed for tiller count in MI (**Fig. 2f**). To investigate why combining SNP and transcriptomic data provides only limited benefit, we identified the genes associated with the SNPs and transcripts used by each model for prediction, defined as those assigned a model importance score greater than zero (**Supplementary Table 2**). We focused on flowering time and biomass as examples because of their agronomic importance and their relatively stronger predictive performance compared to other traits (Figure 2), which provides a more reliable basis for interpreting feature importance scores and SHAP values. We found the highest gene overlap between SNP and transcriptomic models in TX for flowering time (60%), followed by the biomass plasticity model (45%).

All other overlaps were ≤11%, indicating that models predicting the two data types largely rely on distinct sets of genes. Moreover, shared genes had uncorrelated SHAP values (**Supplementary Fig. S2**), consistent with prior findings in maize^11^, suggesting that SNP and transcriptomic models capture complementary information. To evaluate the relative contribution of each data type, we examined feature importance scores from multi-omic models. Transcriptomic features were consistently enriched among predictors with non-zero SHAP values (odds ratios = 5.32–14.8; all FDR-adjusted *p* < 2.2 × 10⁻¹⁶), indicating that transcriptomic data tend to be more informative than SNPs for trait prediction. This finding is consistent with prior results in maize^11^ and highlights the value of transcriptomic data in identifying functionally relevant genes.

To further evaluate the biological relevance of features, we asked whether transcriptomic or SNP data more effectively identify known causal genes. We assessed whether SNP- and transcriptomic-based models assigned non-zero feature importance to benchmark genes, defined as homologs of 1,552 flowering-related genes and 95 biomass-related genes from *Arabidopsis thaliana* and rice (*Oryza sativa*) (**Supplementary Table S3**), as well as 1,245 switchgrass biomass QTL genes from previous studies^6,17,18^ (**Supplementary Table S3; see Methods**). Across feature importance metrics (SHAP and BRR coefficients), transcriptomic models identified more benchmark genes with non-zero importance scores than SNP-based models. For flowering time, 20% of benchmark genes (310 genes) were prioritized by transcriptomic models compared to 8% (129 genes) by SNP models (odds ratio = 2.3, FDR-adjusted *p*=1.8×10⁻^14^). A similar pattern was observed for biomass, with 18% (17 genes) vs. 4% (4 genes) prioritized, respectively (odds ratio = 4.4, FDR-adjusted *p*=0.004; **Supplementary Table S2**). These results suggest that transcriptomic models more effectively identify causative genes, potentially reflecting greater phenotypic influence of gene expression variation compared to SNP variation. It is important to note, however, that gene expression was measured at a single pre-flowering developmental stage. Therefore, the genes identified here reflect transcriptional variation in a specific tissue at that specific time point, and their association with the traits should not be interpreted as evidence that they directly regulate these traits throughout all stages of development.

Having found that transcriptomic models incorporate more benchmark genes for prediction, we next asked which feature importance metric, SHAP or BRR coefficients, more effectively captures these genes in transcriptomic models. Because of the limited number of benchmark genes for biomass (n = 17), we focused this comparison on flowering time. SHAP consistently identified more flowering-related benchmark genes (130 for TX, 105 for MI, and 113 for Diff) than BRR coefficients (77, 54, and 62, respectively), indicating that SHAP provides a more informative measure of feature importance for identifying causal genes. We then evaluated whether benchmark genes were among the top predictors, as would be expected for causal loci. Using the top 5% of features ranked by importance, we observed significant enrichment for benchmark genes only in the TX flowering time model (odds ratio = 3.3, FDR-adjusted *p*=5.2×10⁻⁵). However, non-benchmark genes consistently received higher importance scores than benchmark genes across all models and metrics (FDR-adjusted *p*<2.22×10⁻¹⁶; **Supplementary Fig. S3**). These results suggest that while benchmark genes contribute to prediction, many genes contributing to variation in flowering time and biomass remain uncharacterized.

To identify genes driving model predictions, we examined the most influential genes in the transcriptomic-based models based on SHAP values. For flowering time, 12 genes were consistently ranked in the top 1% across both MI and TX (**Fig. 3a,c**), indicating that they represent a core set of genes that control flowering time across environments. This set included benchmark homologs of *FT* genes (*Pavir.4KG047800*, *Pavir.3KG349500*); genes in this family have been shown to promote flowering in multiple species, including *Arabidopsis* and switchgrass^19^. The core set also included a MADS-box gene (*Pavir.2KG001200*), and a non-benchmark bHLH transcription factor (*Pavir.1NG011300*) homologous to maize and sorghum genes involved in circadian regulation. Despite this shared core set, distinct top predictors emerged in each environment. For instance, the top predictor in TX was a kinesin motor protein (*Pavir.6NG016531*), while in MI, it was a *bZIP* transcription factor (*Pavir.9NG848400*); both are non-benchmark genes from families previously associated with flowering and plant development^20,21^. In the flowering time plasticity model, *Pavir.4KG047800* (*FT*) was the top predictor (**Fig. 3a,d–e**), consistent with its role in environmental responsiveness in *Arabidopsis*^22^. The second-ranked gene, *Pavir.9NG746100*, encodes an AP2/ERF transcription factor; members of this family are known to mediate responses to cold^23,24^ and regulate flowering under variable photoperiods^25^.

**Figure 3.**
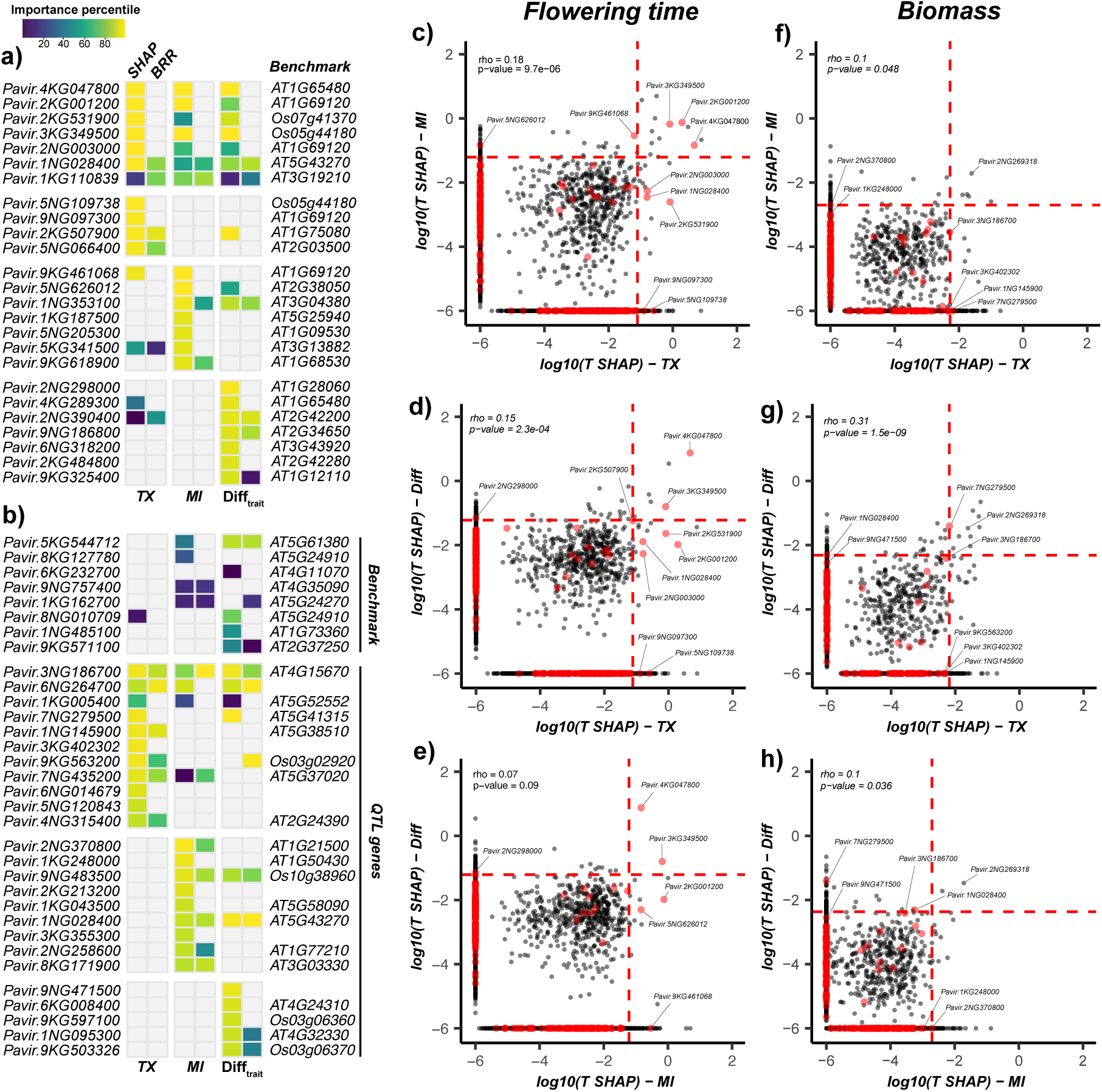
Correlation of transcript importance across MI, TX, and plasticity models for flowering time and biomass. **(a-b)** Feature importance scores for benchmark genes for flowering time (FLT) **(a)** and benchmark genes plus QTL candidate genes for biomass **(b)**. Feature importance was assessed using SHAP values and Bayesian ridge regression (BRR) for two individual environments (TX and MI) and their difference (Diff). Heatmap intensity represents the percentile rank of feature importance within each model, with yellow indicating highest importance. **(c-h)** Comparison of SHAP values between the TX, MI, and Diff models for FL **(c-e)** and biomass **(f-h)**. SHAP values < 10⁻⁶ were set to 10⁻⁶ for visualization. Red dashed lines indicate the 99th percentile of SHAP values. Red features represent benchmark genes **(c-e)** and QTL candidate genes **(f-h)**. Correlations are shown as Spearman rank coefficients (rho).

For biomass, five genes were among the top 1% of important predictors in both MI and TX, four of which encode uncharacterized proteins. The only annotated gene within this top 1% was *Pavir.2NG269318,* a meso-2,6-diaminoheptanedioate carboxy-lyase involved in lysine biosynthesis (**Fig. 3f**). In Arabidopsis, lysine biosynthesis mutants show reduced photosynthesis and growth^26^, supporting a functional link to biomass. This gene also ranked highly in the biomass plasticity model (**Fig. 3g–h**). Environment-specific predictors included a kinesin motor protein (*Pavir.6NG016531*) in TX and a glycosyltransferase (*Pavir.1KG124131*) in MI; overexpression of glycosyltransferases has been linked to biomass increases in *Arabidopsis*^27^. The top predictor in the biomass plasticity model was an AP2/ERF transcription factor (*Pavir.7NG295800)*, previously associated with biomass in switchgrass^6^. As in flowering time, AP2/ERFs emerged as regulators of G×E responses, consistent with their roles in mediating plant responses to climatic and photoperiod variation^23–25^. Taken together, these results demonstrate that transcriptomic models capture both shared and environment-specific biological signals, effectively identifying benchmark genes and uncovering novel, uncharacterized candidates that may underlie natural variation in traits and their plasticity.

### Gene effects are genotype and environment dependent

Building on our finding that models recover both benchmark and novel candidate genes, we next asked whether gene effects vary among genotypes across environments. To quantify genotype-specific contributions of individual genes, we computed local SHAP values for each genotype, where positive SHAP values indicate that higher feature values (e.g., gene expression) are associated with higher trait values, and negative values indicate the opposite. We focused on the top 20 genes predictive of flowering time and biomass, and used SHAP profiles to cluster genotypes based on shared patterns of gene importance. In each data set, these clusters showed distinct trait distributions (**Supplementary Fig. S4, S5**), demonstrating that SHAP profiles reflect transcriptomic influences on trait variation and capture genotype-level differences in how genes contribute to trait expression.

To investigate G×E interactions, we analyzed transcriptome-based models predicting trait value differences across environments. Clustering genotypes based on their local SHAP profiles revealed nine distinct groups with varying flowering time plasticity (**Fig. 4a**). In these models, positive SHAP values indicate the contribution of predictors to delayed flowering time in TX relative to MI, while negative values indicate earlier flowering time in TX. Clustering was primarily driven by *Pavir.4KG047800* (FT homolog) and *Pavir.9NG746100* (AP2/ERF transcription factor). Genotypes in clusters 1 and 2 exhibited positive SHAP values, corresponding to delayed flowering time in TX. To understand this variation, we examined gene expression differences across environments. Genotypes with higher FT and AP2/ERF expression in TX tended to flower earlier in TX compared to MI, while genotypes with stable expression tended to flower later in TX **(Fig. 4b–c)**. These results suggest that expression plasticity of FT and AP2/ERF contributes to adaptive shifts in flowering time. Consistent with this, FT and AP2/ERF genes are known regulators of flowering and environmental responsiveness across species^2,28–31^.

**Figure 4.**
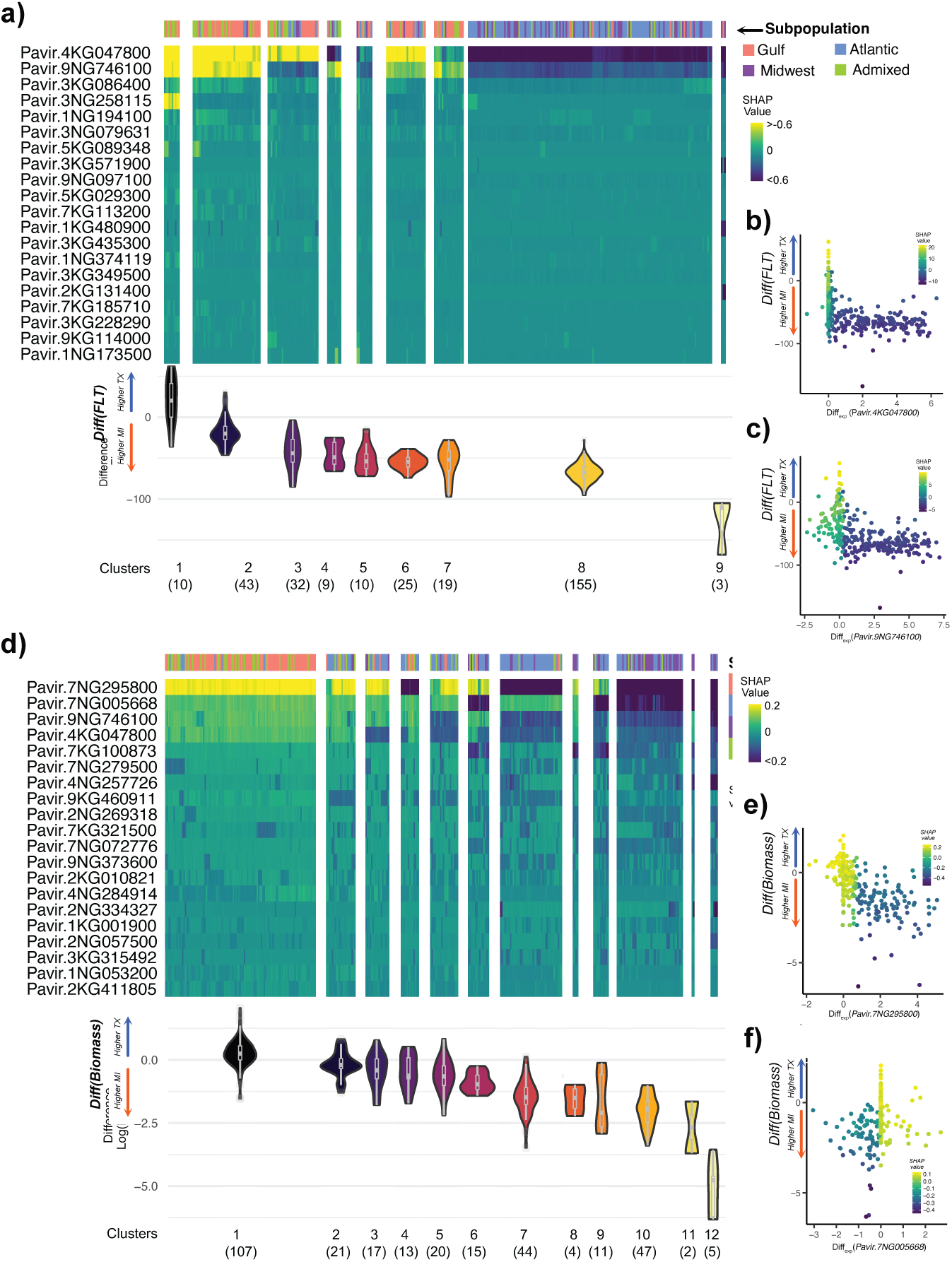
Genotype-dependent effects of top transcriptomic predictors of trait plasticity. **(a, d)** Euclidean clustering of SHAP values for the top 20 genes (y-axis) across genotypes (x-axis) for flowering time (FLT) **(a)** and biomass **(d)** plasticity models. For visualization, SHAP values were clipped to ±0.6 for flowering time and ±0.2 for biomass **(b–c, e–f)** Relationship between gene expression plasticity (difference in expression level [TPM] between TX and MI; x-axis) and trait plasticity (difference in trait value between TX and MI; y-axis) for the top two predictors of FLT **(b–c)** and biomass **(e–f)**. Colors indicate the SHAP values.

In the biomass model, SHAP values grouped genotypes into 12 distinct clusters (Fig. 4d). The top predictors included two AP2/ERF transcription factors (Pavir.7NG295800 and Pavir.9NG746100, ranked 1 and 3), a gene of unknown function (Pavir.7NG005668, rank 2), and FT (Pavir.4KG047800, rank 4). Notably, both Pavir.9NG746100 and FT were also among the top predictors for FLT, reinforcing their roles in regulating G×E responses across traits^30^. Clustering revealed distinct biomass patterns. Cluster 1 included genotypes with higher biomass in TX compared to MI, genotypes in clusters 2–5 showed similar biomass across environments, and those in clusters 6–12 had higher biomass in MI compared to TX (**Fig. 4d**). The top predictors exhibited opposing SHAP value directions across clusters (e.g., clusters 3–7, 9, 11), indicating decoupled gene effects. For instance, genotypes with higher expression of *Pavir.7NG295800* in TX had negative SHAP values (i.e., higher biomass in MI; **Fig. 4e**), while *Pavir.7NG005668* showed the opposite pattern (**Fig. 4f**). Similar genotype-specific contrasting SHAP values were observed for the top 20 predictors in single-environment models (**Supplementary Fig. S4, S5**), highlighting the genotype dependent effect of T features. Together, these findings demonstrate that gene effects on flowering time and biomass are both genotype and environment dependent and that variation in gene expression contributes to trait plasticity across environments.

### Gene-by-gene interactions reveal environment-specific regulatory architecture

In previous sections, we showed that gene effects on traits are both genotype- and environment-dependent. Because each genotype carries a unique combination of alleles, we hypothesized that variation in gene composition among genotypes could contribute to differences in gene–gene interactions, ultimately influencing trait expression. To evaluate feature interactions and how they vary across environments, we used SHAP interaction scores^32^, which quantify pairwise feature interactions and may reflect epistatic relationships within the predictive model. For interaction analyses, we focused on transcriptomic models because transcript-based predictors were enriched among the most important features in multi-omic models and identified a greater number of benchmark genes than SNP-based models. In addition, gene expression data provide a more biologically interpretable framework for assessing Gene × Gene interactions, as transcripts more directly reflect gene activity and regulatory processes. We first asked whether the number and magnitude of interactions were consistent across models. Genes important for flowering time showed both stronger and more interactions than those for biomass (**Fig. 5a–b**). This likely reflects the underlying genetic architecture of flowering time, which is regulated by central hubs with large effect sizes^33,34^, in contrast to the more diffuse, polygenic control of biomass. Such large-effect loci may enhance the detectability of interaction signals.

**Figure 5.**
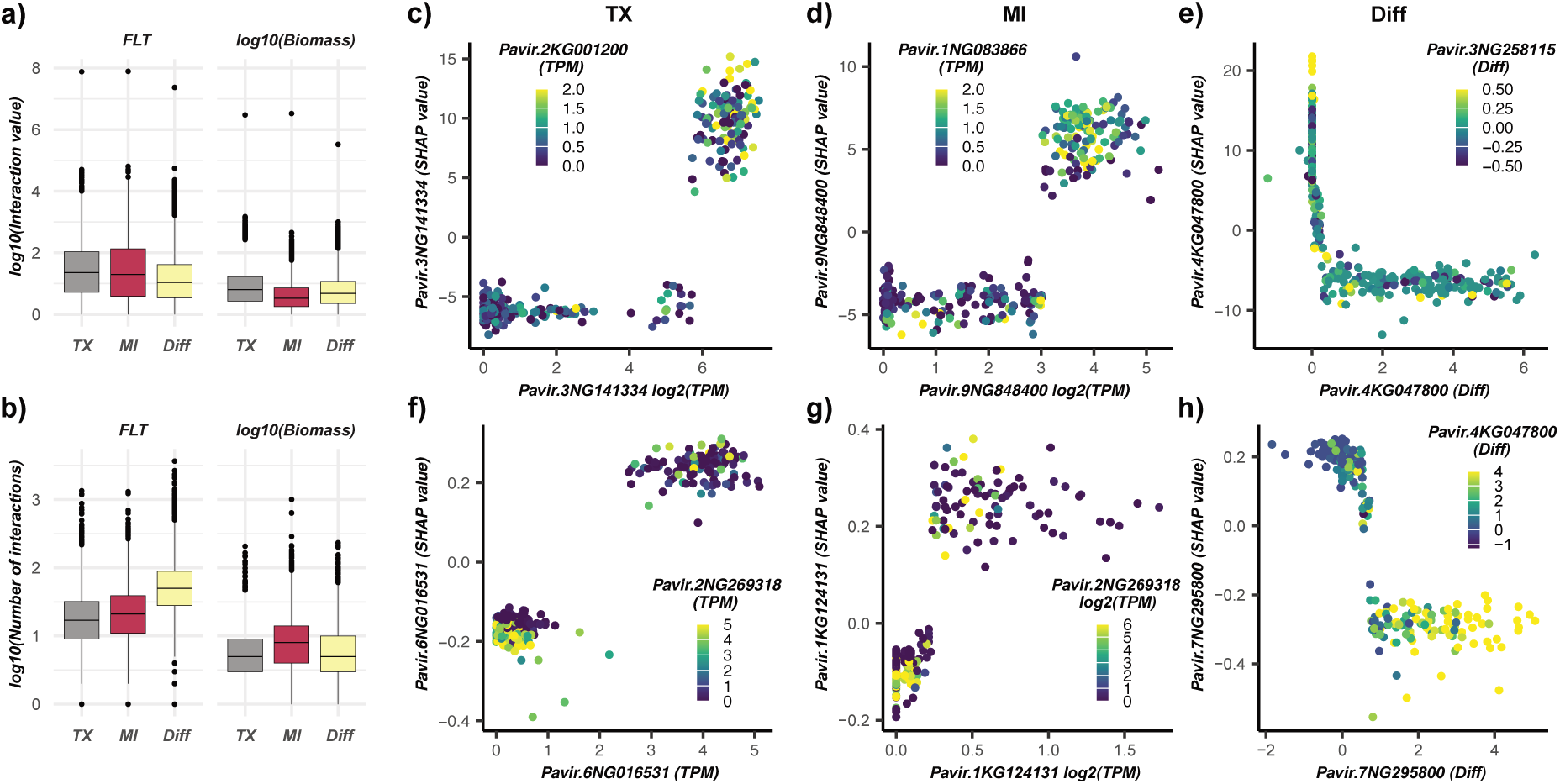
Gene–gene interaction profiles associated with flowering time and biomass. **(a)** Distributions of SHAP interaction values from transcriptomic models for the flowering time (FLT; left) and biomass (right) models. **(b)** Number of gene–gene interactions per gene for FLT (left) and biomass (right). **(c-d)** Examples of top interactions for FLT **(c)** and biomass **(d)**.

Genes with strong interactions can modulate the effects of other genes and contribute to non-additive genetic variance. To identify hub genes, we ranked the top 100 genes by the number of SHAP interactions. In flowering time models, two FT homologs (*Pavir.4KG047800*, *Pavir.3KG349500*) and two MADS-box genes consistently ranked among the top interactors in both MI and TX, each with over 169 interactions (**Supplementary Table S5**). Additionally, two non-benchmark genes ranked first in MI (*Pavir.9NG848400*, a bZIP transcription factor) and TX (*Pavir.3NG141334*, an FKBP). Both gene families have previously been implicated in flowering regulation^21,35^, supporting their potential roles as flowering time hub genes. In the plasticity model, *Pavir.4KG047800* (FT) had the highest interaction count (3,670), reinforcing the hypothesis that it plays a central role in mediating G×E responses. Other top benchmark genes included BES/BZR, AGC kinase, and bHLH transcription factors, all known regulators of environmental responses^36–38^. Top non-benchmark interactors included a cytochrome P450 and an AP2-like transcription factor, members of gene families linked to stress adaptation and plasticity^39,40^. Together, these results demonstrate that transcriptomic models capture both canonical flowering regulators and novel candidate co-regulators of flowering time and its plasticity.

In contrast, for biomass models, no benchmark genes were among the top 100 SHAP interactors; instead, several non-benchmark genes with potential roles in biomass-related traits were identified (**Supplementary Table S6**). For example, in TX, *Pavir.5NG530000* (*ULTRAPETALA* homolog) ranked second (166), and in MI, the top interactor was *Pavir.1KG124131* (*Glycosyltransferase*; 1005). *ULTRAPETALA* genes have been linked to meristem size and plant architecture^41^, while glycosyltransferases have been associated with biomass accumulation^27^. In the plasticity model, the top interactor was *Pavir.7NG295800* (AP2/ERF; 233). Notably, the FT homolog *Pavir.4KG047800* ranked 11th (105), highlighting the potential role of the vegetative-to-reproductive transition in biomass accumulation and its contribution to plasticity.

Examining interactions in detail, we found that *Pavir.4KG047800* (*FT* homolog) and *Pavir.2KG001200* (*MADS-box AP1 homolog*) were co-expressed in both MI and TX, where high expression correlated with earlier flowering (**Supplementary Fig. S6a–b**), consistent with FT’s activation of *AP1*^42^. In the plasticity model, interactions between two FT homologs (*Pavir.4KG047800* and *Pavir.3KG349500*) showed environment-dependent *effects*; genotypes that failed to upregulate FT-like genes in Texas flowered later, underscoring FT’s central role in flowering plasticity (**Supplementary Fig. S6c**). We also identified a non-benchmark gene, *Pavir.3NG141334* (an FKBP-like gene), which interacts with *AP1* and is associated with delayed flowering in TX when both genes show high expression (**Fig. 5c**), paralleling *FKBP12–CONSTANS* interactions in Arabidopsis^35^. In MI, we identified an interaction between a *bZIP* (*Pavir.9NG848400*) and a glucosyltransferase (*Pavir.1NG083866*) (**Fig. 5d**). Members of both genes families have established roles in *FT* activation complexes and hormonal regulation of flowering^42,43^. Another glucosyltransferase (*Pavir.3NG258115*) interacted with *Pavir.4KG047800* (*FT*) in the plasticity model, where genotypes with stable *FT* and high *Pavir.3NG258115* expression flowered later in TX (**Fig. 5e**).

SHAP interaction values revealed that interactions between Pavir.2NG269318 (carboxy-lyase) and Pavir.6NG016531 (kinesin motor protein) in Texas, and between Pavir.2NG269318 and Pavir.1KG124131 (glycosyltransferase) in Michigan, contributed to model predictions of biomass **(Fig. 5f–g)**. In both cases, high *Pavir.2NG269318* expression was associated with negative SHAP values for *Pavir.6NG016531* and *Pavir.1KG124131*, indicating that under high *Pavir.2NG269318* expression, these growth-related genes contribute negatively to biomass. One potential explanation for these interactions is that high Pavir.2NG269318 expression alters metabolic flux in a way that limits the effectiveness of cell expansion, mediated by kinesin motor proteins^44^, and cell-wall biosynthesis, driven by glycosyltransferases^45^. In the plasticity model, *Pavir.7NG295800* (AP2/ERF) interacted with *Pavir.4KG047800* (*FT*), where higher expression in TX relative to MI for both genes corresponded to reduced biomass in TX compared to MI. This pattern may indicate that elevated expression of *FT* (a flowering regulator) and an AP2/ERF transcription factor (a regulator of growth and development)^30^ shortens the vegetative phase in TX, resulting in lower biomass accumulation. Together, these results suggest that local interpretation can recover known flowering regulators and highlight novel, environment-specific gene relationships, offering a framework for generating hypotheses about the molecular basis of flowering, biomass accumulation, and G×E plasticity.

## Conclusion

By integrating genomic and transcriptomic data with interpretable machine learning, we show that (i) information from models based on transcripts contributing environment-sensitive signals doesn’t overlap with that captured by models using genetic variance; (ii) gene effects are strongly genotype- and environment-dependent; and (iii) local interpretation reveals biologically meaningful, environment-specific gene–gene interactions. FT homologs consistently emerged as central hubs for flowering time and, to a lesser extent, biomass, highlighting the pleiotropic role of flowering regulators in connecting reproductive transitions with vegetative growth. These findings translate complex model predictions into mechanistic hypotheses about causal genes and interactions underlying trait variation within and across environments. Gene expression was measured at a single developmental stage and from a single sample per genotype– environment combination; this experimental design prioritized broad genetic and environmental sampling across a large diversity panel. Consequently, the candidate genes and interactions identified should be viewed as hypothetical associations rather than definitive causal claims. Together, our results prioritize candidate regulators for experimental validation and provide a general framework for dissecting the genetic basis of complex traits and their plasticity in plants.

## Materials and Methods

### Phenotypic data

The diversity panel used in this study were previously described^6^. Briefly, the switchgrass diversity panel was established in 2018, with plants arranged in a honeycomb design such that each individual had four nearest neighbors located 1.5 m away across 10 sites in the United States. The locations in Michigan and Texas were chosen for the current study because of their contrasting environmental conditions and latitudes, which have been observed to influence important trait variation in switchgrass. In Michigan, the diversity panel was grown at the Michigan State University Kellogg Biological Station (KBSM) (latitude: 42.419618, longitude: – 85.371266, elevation: 289 meters). In Texas, plants were grown at the University of Texas J.J. Pickle Research Campus (latitude: 30.383979, longitude: –97.729383, elevation: 235 meters). Genotypes were replicated established as a single-block planting at each location, with genotypes distributed throughout the field. Because only a single replicate per genotype was established at each site, formal spatial correction was not performed, as genotype and within-site spatial effects could not be estimated independently. However, the AP13 reference genotype was distributed throughout the experimental plots and did not show evidence of substantial spatial heterogeneity at the Michigan site (data not shown).

In total, six traits were measured in 2021 for a subset of 426 genotypes: green-up (GR), emergence (EM), flowering time (FLT), tiller count (TC), height of the panicle (HT), and biomass. GR was measured as the day of the year when plants began to green up. EM was measured as the day of the year when panicles began to emerge. FLT was recorded as the day of the year when flowering started. TC and HT were measured at the end of the season, with height recorded as the mean panicle apex in centimeters. For biomass, whole plants were harvested and weighed (recorded in grams), and dry weight was estimated for each sample based on a ∼500 g subsample of fresh material. The estimated dry biomasses were log-transformed to normalize the data. Phenotypic data and correlations across locations are presented in **Supplementary Figure 1.**

To partition phenotypic variance into genetic (G), environmental (E), and genotype-by-environment interaction (G×E) components, we fit the following linear mixed model using the sommer package^46^ in R:

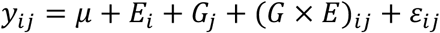

where *y_i_*_j_ represents the phenotypic value of genotype *j* in environment *i*, *μ* is the overall mean, *E_i_*is the random effect of environment, *G*_j_is the random genetic effect modeled using the genomic relationship matrix (kinship), (*G* × *E*)*_i_*_j_ represents the genotype-by-environment interaction effect, and *ε_i_*_j_ is the residual error. Variance components were extracted from the fitted model, and the proportion of total phenotypic variance explained by each component was calculated.

### Whole genome data and genetic variants

Whole-genome sequencing data from the switchgrass diversity panel, comprising 726 genotypes, were downloaded from the NCBI SRA under BioProject PRJNA622568. The genotypes were sequenced using Illumina HiSeq X10 and Illumina NovaSeq 6000 paired-end sequencing (2 × 150 bp) at the HudsonAlpha Institute for Biotechnology and the Joint Genome Institute^6^. Genetic variant detection followed the GATK Best Practices (Broad Institute). Briefly, raw reads were first trimmed as paired-end reads using fastp with default parameters^47^. Clean reads were then mapped to the AP13 HAP1 v6.1 genome (used with permission) using BWA-MEM^48^ with default parameters. The resulting BAM files were sorted using samtools^49^. Next, duplicate reads were marked using picard toolkit MarkDuplicates. Variant calling was performed using GATK HaplotypeCaller, generating a GVCF (genomic variant calling file) for each sample. The GATK GenomicsDBImport function was then used to merge GVCFs from all 732 samples into a GenomicsDB. Finally, GATK GenotypeGVCFs was used to identify genetic variants in each sample. The resulting VCF file was filtered using BCFtools to retain the 426 genotypes for which phenotypic data were collected, as well as SNPs with no missing data and a minor allele frequency (MAF) > 0.05. To reduce redundancy among highly correlated markers, linkage disequilibrium (LD) pruning was performed using the +prune plugin in BCFtools, with pairwise LD measured as the squared correlation coefficient (r²). SNPs with r² > 0.1 within a 5 Mb sliding window were removed. The final VCF file contained 220,263 SNPs.

### Leaf collections and RNA extractions

Leaves of mature plants for each of the 426 genotypes were collected from the field in both in TX and MI. We timed collections at both sites to best approximate the same pre-flowering developmental stage. Leaf collections at the common garden located at J.J. Pickle Research Campus in Austin, TX were conducted on May 5 and May 6, 2021. Leaf collections from the common garden at the Kellogg Biological Station in Hickory Corners, MI were conducted on June 22^nd^ and June 24^th,^ 2021. We collected the highest fully expanded leaves on tillers at both locations between 10 am and noon. For each genotype, the proximal ∼4cm of a collected leaf was placed in a 2mL microcentrifuge tube and flash frozen in liquid nitrogen. Tissue was then transported to -80 freezers at The University of Texas and Michigan State University on dry ice.

For RNA extraction, approximately 200 mg of leaf tissue was homogenized into a fine powder in 2.0-mL Premium Microcentrifuge Tubes (Fisher Scientific) using stainless steel beads on a Geno/Grinder 2000 (Spex SamplePrep). The tubes were kept in a pre-frozen microcentrifuge tube rack under liquid nitrogen to maintain the samples in a frozen state during homogenization. Total RNA was isolated using the Direct-zol™ RNA Miniprep Kit (Zymo Research) following the manufacturer’s instructions. In brief, 1 mL of TRIzol™ Reagent (Invitrogen) was added immediately to the frozen powdered tissue, followed by vigorous vortexing to fully resuspend the homogenate. The lysate was centrifuged at 12,000 × g for 5 min at 4°C to remove insoluble debris, and 400 µL of the clear supernatant was transferred to a new RNase-free microcentrifuge tube. An equal volume (400 µL) of 100% ethanol was added to the supernatant and mixed thoroughly. The mixture was then loaded onto a Direct-zol™ spin column, and RNA purification and on-column DNase I treatment were performed according to the manufacturer’s protocol. RNA was eluted by adding 40 µL of DNase/RNase-free water directly to the column matrix and centrifuging. The integrity and concentration of the RNA preparations were checked initially using Nano-Drop (Nano-Drop Technologies) and then by BioAnalyzer (Agilent Technologies).

### RNA-seq data normalization

RNA-seq was performed using NovaSeq 6000 paired-end sequencing (2 x 150 bp) at the Joint Genome Institute. Raw fastq file reads were filtered and trimmed using the JGI QC pipeline. Raw reads were evaluated for artifact sequence by kmer matching (kmer=25), allowing 1 mismatch, and detected artifacts were trimmed from the 3’ end of the reads using BBDuk, a tool included in the BBMap package (https://sourceforge.net/projects/bbmap/). RNA spike-in reads, PhiX reads and reads containing any Ns were removed. Quality trimming was performed using the phred trimming method set at Q6. Finally, reads under 50 bases in length were removed. Filtered reads from each library were aligned to the AP13 HAP1 v6.1 genome using HISAT2 version 2.2.0^50^. Strand-specific coverage bigWig files (fwd and rev) were generated using deepTools v3.1^51^. featureCounts^52^ was used to generate the raw gene counts file using AP13 HAP1 v6.1 gff3 annotations. Only primary hits assigned to the reverse strand were included in the raw gene counts. For downstream machine learning analysis, raw read counts for each gene were divided by the gene’s length (in kilobases) to calculate Reads Per Kilobase (RPK). Then, RPK values for all genes within a sample were summed, and each RPK value was divided by this sum to obtain Transcripts Per Million (TPM) values. To remove genes with zero or nearly zero variance (>95% of samples sharing the same transcript level), we used the nearZeroVar function from the R caret package. After filtering, TPM counts for 49,985 genes were retained in the final dataset. Before further analysis, TPM values were transformed (log₁₀(TPM + 1)).

### Correlations between genetic relationships, gene expression, and phenotypic variation across environments

The genomic relationship matrix (G) was estimated using centered and standardized SNP information, following the formula G = *ZZ*′/*p* ^53^, where *p* represents the number of SNPs and *Z* is the matrix of centered and standardized SNPs. For gene expression, Pearson correlations of TPM values between genotypes were computed using the cor.test function in R, generating an expression correlation (eCor) matrix for each environment. For phenotypic data, the Euclidean distance between genotypes was calculated for each trait. The correlation between the G, eCor, and phenotypic distance was assessed using Pearson correlation. Additionally, genomic heritability (ℎ^2^), defined as the proportion of variance in a trait that can be explained by a linear regression on markers^54^, was estimated for all traits.

### Predictive modeling and feature importance

Because of the well-known relationship between subpopulations and phenotypic variation in switchgrass, a baseline for the prediction models was established using the first five principal components (PCs) derived from SNPs to predict phenotypic values. The first five PCs were chosen as a proxy for population structure because, beyond the fifth PC, the proportion of variance explained is lower than 1% (Supplementary Figure 2). Genotypes for the training and test sets were selected using stratified sampling with ten equally spaced quantile bins (10% intervals from 0 to 1), ensuring proportional sampling of individuals in both sets from each quantile. To maintain consistency across traits and environments, this stratification approach was applied to log-transformed biomass in TX, ensuring that the same individuals were used in the training and test sets for all traits and environments.

For each trait, predictions were performed using a linear model, Bayesian Linear Regression (BRR), and a non-linear model, eXtreme Gradient Boosting (XGBoost). The BRR models were implemented in R using the BGLR package^55^ with 12,000 iterations, where the first 2,000 iterations were discarded as burn-in. XGBoost was implemented in Python using the Scikit-Learn package^56^. To optimize the hyperparameters of the XGBoost models, a Bayesian optimization using the Tree-structured Parzen Estimator (TPE) algorithm was implemented using the HyperOpt package^57^. The hyperparameter search space was defined as follows: the learning rate was sampled from a uniform distribution between 0.001 and 0.3; the maximum tree depth was chosen from discrete values (3, 5, 10); the subsample ratio was sampled from a uniform distribution between 0.8 and 0.9; feature subsampling was sampled from a uniform distribution between 0.5 and 1.0; and the number of estimators was chosen from discrete values (50, 100, 150, 200). The optimization process was assessed using R² through 5-fold cross-validation. Note that Figure 2 includes only model performance on the testing dataset. Cross-validation was performed within the training set for hyperparameter tuning. Once the optimal parameters were identified, the model was retrained on the full training set and used to predict the testing dataset. Cross-validation R² values are provided in Supplementary Table 1. The search process was executed for 100 iterations, after which the best-performing hyperparameters were selected. A final model was then built using all genotypes in the training set and applied to the test set to evaluate performance using R².

To evaluate the global contribution of features to model predictions, feature importance values were obtained from both the BRR and XGBoost models. For BRR, feature importance was measured as the estimated posterior mean of feature effects, calculated using the R package BGLR^55^, with the absolute values of these effects used for interpretation. For XGBoost, feature importance was assessed using gain-based importance, extracted from the fitted models using the “feature_importances_” function from the Scikit-Learn package^56^. To reduce variability in feature importance estimates caused by stochastic elements in the boosting process, such as random subsampling and feature selection, the importance scores were averaged across 10 model runs using the same set of optimized hyperparameters and training dataset. To assess the local contribution of features to individual predictions (i.e., genotypes), SHapley Additive Explanations (SHAP) values^58^ were obtained from XGBoost. SHAP values were computed in each of the 10 runs using the “explainer_model” function from the SHAP package^58^, and were then averaged across runs. SHAP values are expressed in the same units as the predicted phenotype and represent the additive contribution of each feature to individual predictions.

### Feature selection and feature interactions

Feature selection was initially performed using XGBoost feature importance scores. To reduce the influence of stochastic variation, importance values were averaged across 10 independent model runs, and only features with non-zero average importance were retained. Retained features were ranked according to their mean importance score, and nested subsets containing the top 100, 90, 80, 70, 60, 50, 40, 30, 20, 10, 8, 6, 4, 2, and 1% of features were evaluated. New models were trained for each subset using the same hyperparameter optimization and cross-validation procedures described above (Supplementary Table 1), and the subset producing the highest cross-validated R² was selected. To provide an interpretable assessment of predictor importance, SHAP values were calculated for the final models, and global feature importance was estimated as the mean absolute SHAP value across all models runs. Feature interactions were quantified using the shap_interaction_values function implemented in the SHAP package^58^. Due to the high computational cost of estimating SHAP interaction values, interaction analyses were performed only for the top 5,000 features with the highest global SHAP importance in the final models.

### Identification of candidate genes for biomass

For flowering time, we downloaded 687 and 157 benchmark flowering-time genes (**Supplementary table 3**) from the TAIR and FLOR-ID databases, respectively; these genes are known to regulate flowering in *Arabidopsis*. We also included 237 switchgrass genes associated with flowering time and homologous to 130 *Arabidopsis* and 107 rice genes previously identified in the literature and databases as flowering-time genes^59,60^, resulting in a total of 1,081 genes.

For biomass, we downloaded 63 benchmark genes from TAIR (**Supplementary table 3**). Because few experimentally validated biomass genes are available, we also compiled 1,896 candidate genes from previous studies: 32 from QTL analyses^17,18^ and 1,864 from GWAS analyses^6^. These candidates were originally identified using earlier versions of the switchgrass reference genome (v1.1, v3.1, v4.1, and v5.1). To map them to the most recent reference genome (AP13 v6.1, Hap1), we used OrthoFinder and GENESPACE with default parameters, restricting homolog identification to syntenic orthologs to ensure reliable cross-version mapping (**Supplementary table 4**).

## Data availability

The raw RNA sequencing data for all libraries are available under BioProject PRJNA1402996 at https://www.ncbi.nlm.nih.gov/bioproject/PRJNA1402996. The TPM count matrix used In this study is available in the Zenodo repository at https://zenodo.org/records/21781035.

## Code availability

Custom pipelines for SNP calling, kinship, expression correlation matrices, heritability, predictive modeling, and identification of syntenic orthologous genes are available at https://github.com/pauloizquierdo/switchgrass-GxE-modeling.git.

## Acknowledgments

We thank the field technicians and volunteers who maintained field sites and assisted with phenotype and leaf tissue collections, including L. Vormwald, M. Ryskamp, P. Wang, N. Emery, A. VanWallendael, T. Zambiasi, K. Segura Abá, T. Ranaweera, J. Yin and C. Malmstrom. We thank the Department of Energy Joint Genome Institute and collaborators for pre-publication access to the genome of *Panicum virgatum* AP13 HAP1 v6.1 for RNA-seq analysis. This material is based upon work supported by the Great Lakes Bioenergy Research Center, U.S. Department of Energy, Office of Science, Biological and Environmental Research Program under Award Numbers DE-SC0018409 and DE-FC02-07ER64494. This work was also supported by the U.S. Department of Energy, Office of Science, Biological and Environmental Research Program under Award Number DE-SC0021126. Support was further provided by the National Science Foundation Long-Term Ecological Research Program (DEB-1832042) at Kellogg Biological Station. Work conducted by the DOE Joint Genome Institute (proposal 10.46936/10.25585/60000516), a DOE Office of Science User Facility, is supported by the Office of Science of the U.S. Department of Energy under Contract No. DE-AC02-05CH11231.

## Author Contributions

P.I., D.L., T.J., M.L. and S.H.S. designed research. P.I., D.L., T.J., X.W., Y.Y., C.D., A.L., J.B. and S.H.S. performed research. P.I. and S.H.S analyzed data; and P.I., D.L., T.J., M.L., and S.H.S wrote the paper. All authors read and approved the final manuscript.

## Supplementary figures

**Supplementary Figure 1.**
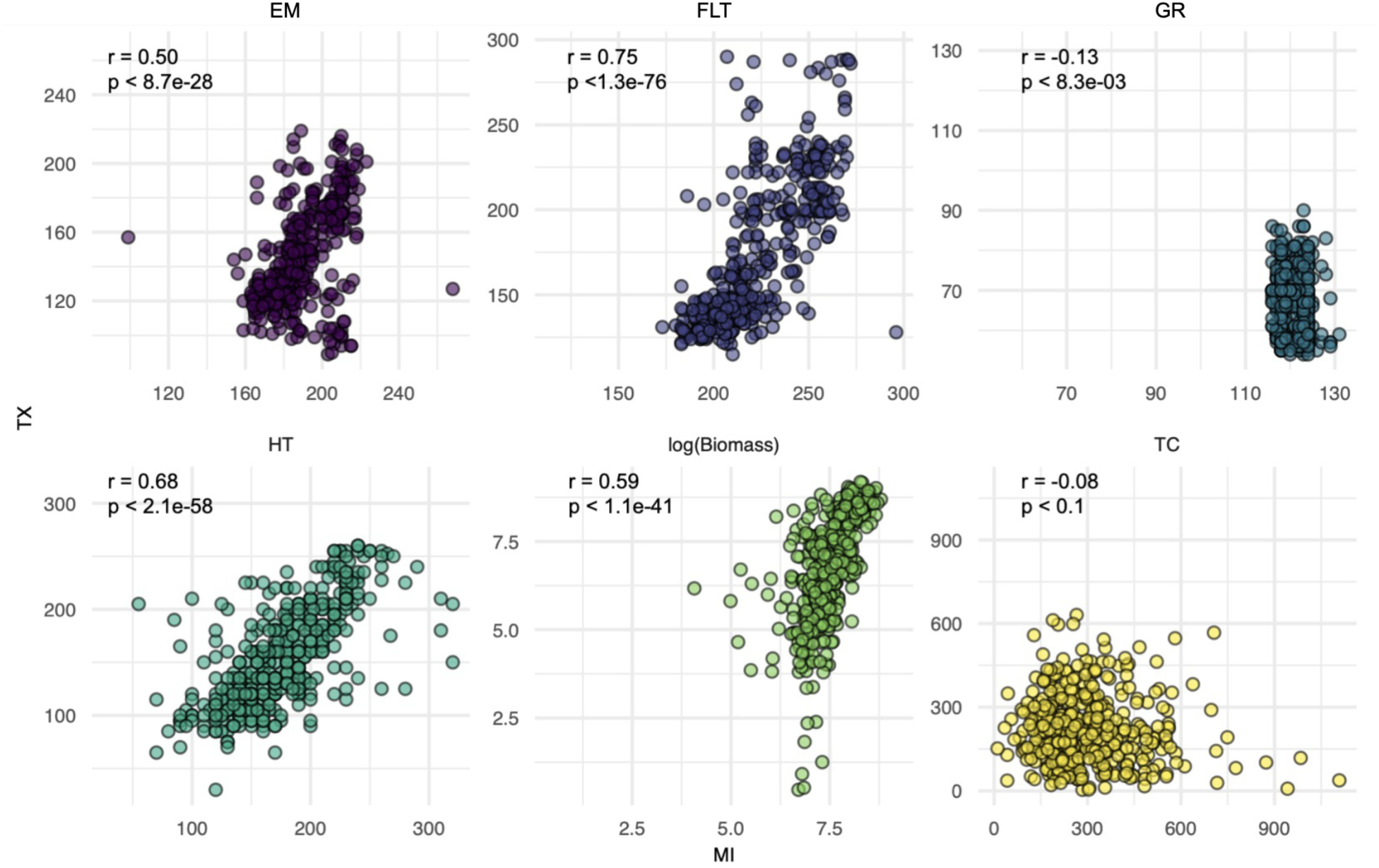
Pearson correlation (r) between traits across environments (x-axis: MI; y-axis: TX) for green-up (GR), emergence (EM), flowering time (FLT), tiller count (TC), panicle height (HT), and log-transformed biomass (measured in g).

**Supplementary Figure 2.**
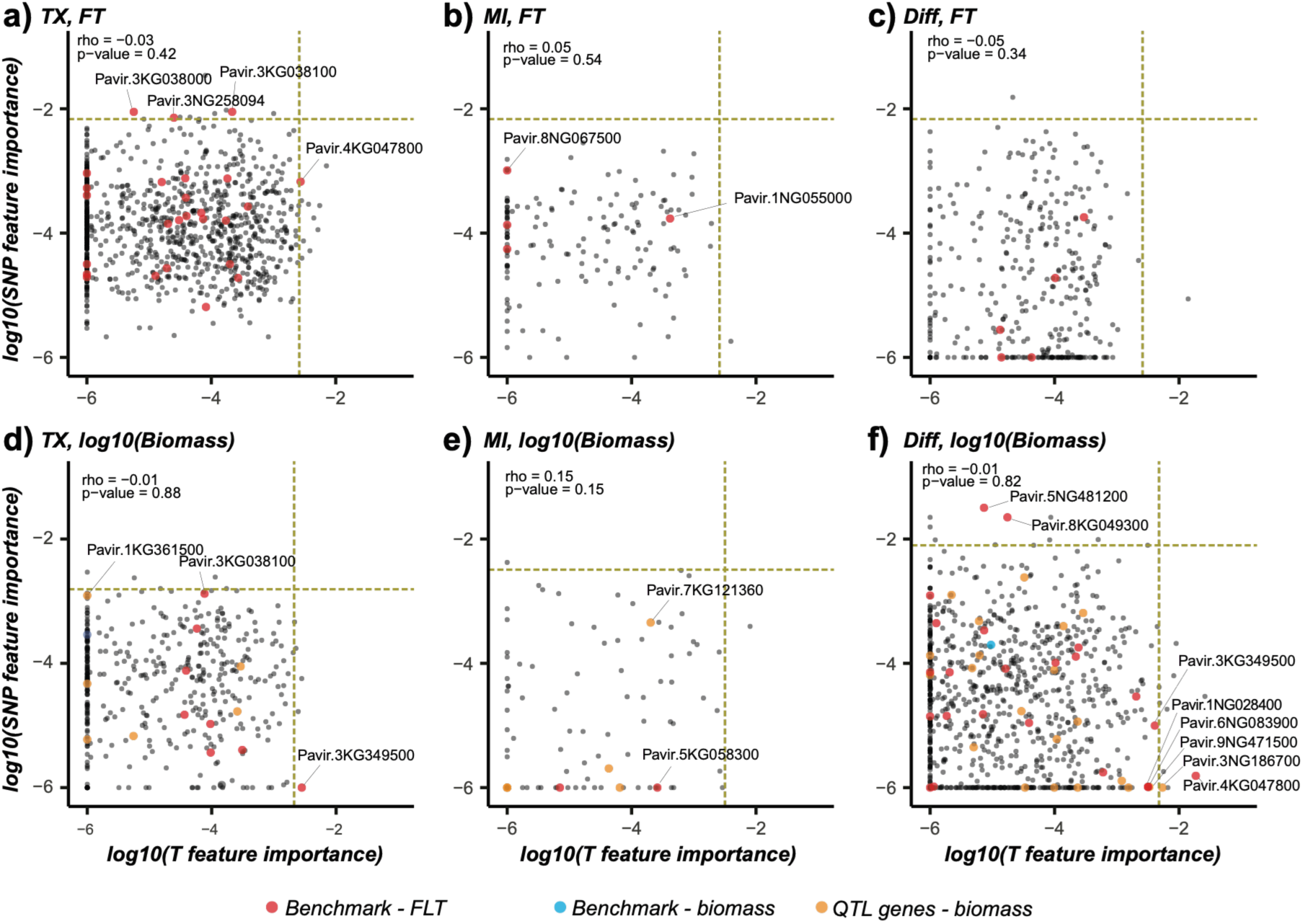
Density scatterplots of importance scores for SNPs (y-axis) and transcripts (x-axis) for SNP:T pairs associated with flowering time (FLT) (a–c) and biomass (d–f) in TX, MI, and difference models. Thresholds (yellow dotted lines) represent the 99th percentile of importance scores. Spearman’s rank correlation (ρ) quantifies correspondence between SNP and transcript importance.

**Supplementary Figure 3.**
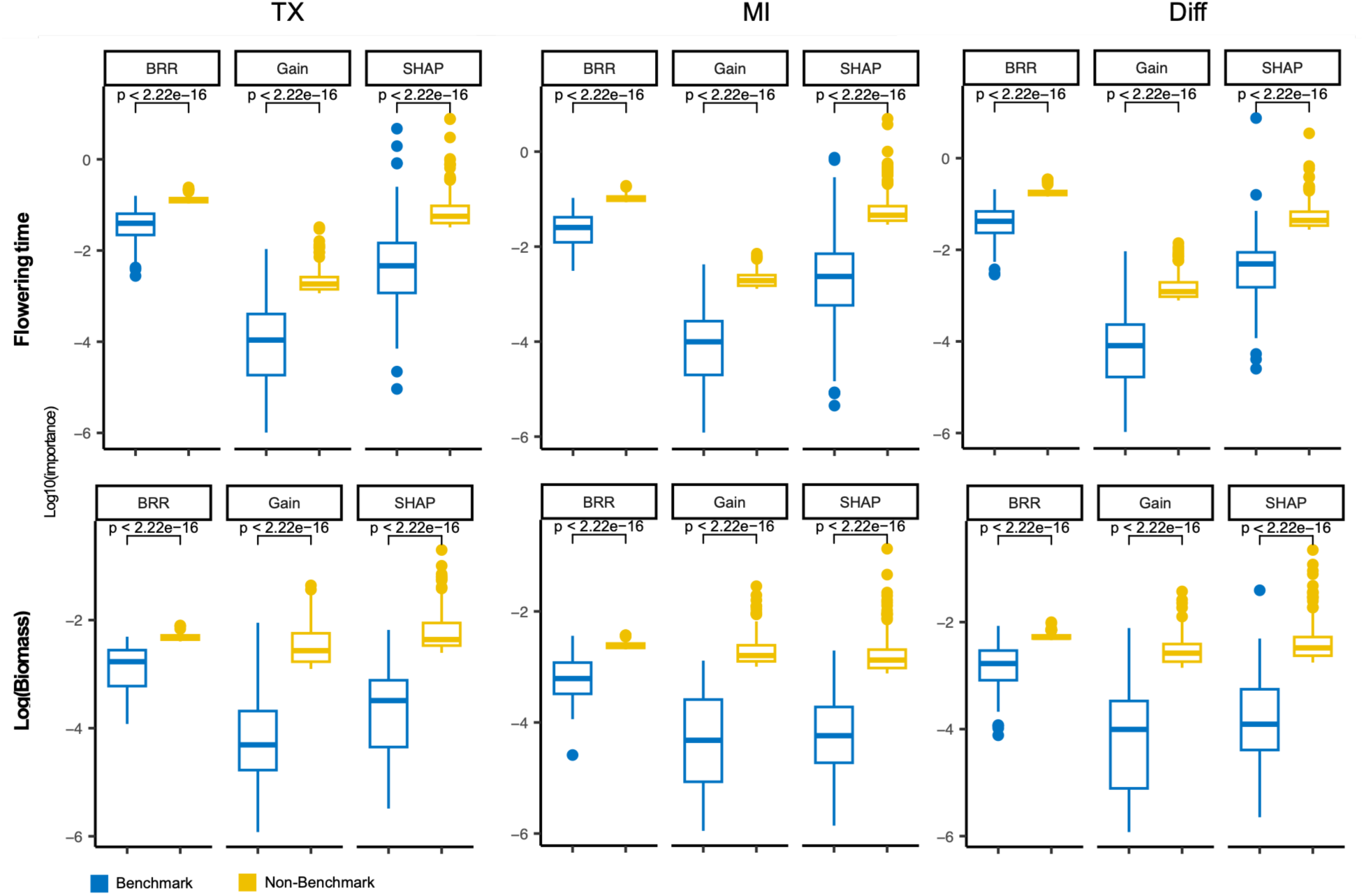
Comparison of transcriptomic importance scores between benchmark and non-benchmark genes associated with flowering time and biomass. P-values were calculated using two-sided Wilcoxon rank-sum tests and adjusted for multiple testing using the Benjamini–Hochberg false discovery rate (FDR). BRR, Bayesian ridge regression coefficients; Gain, XGBoost gain importance; and SHAP, SHapley Additive exPlanations values derived from XGBoost models

**Supplementary Figure 4.**
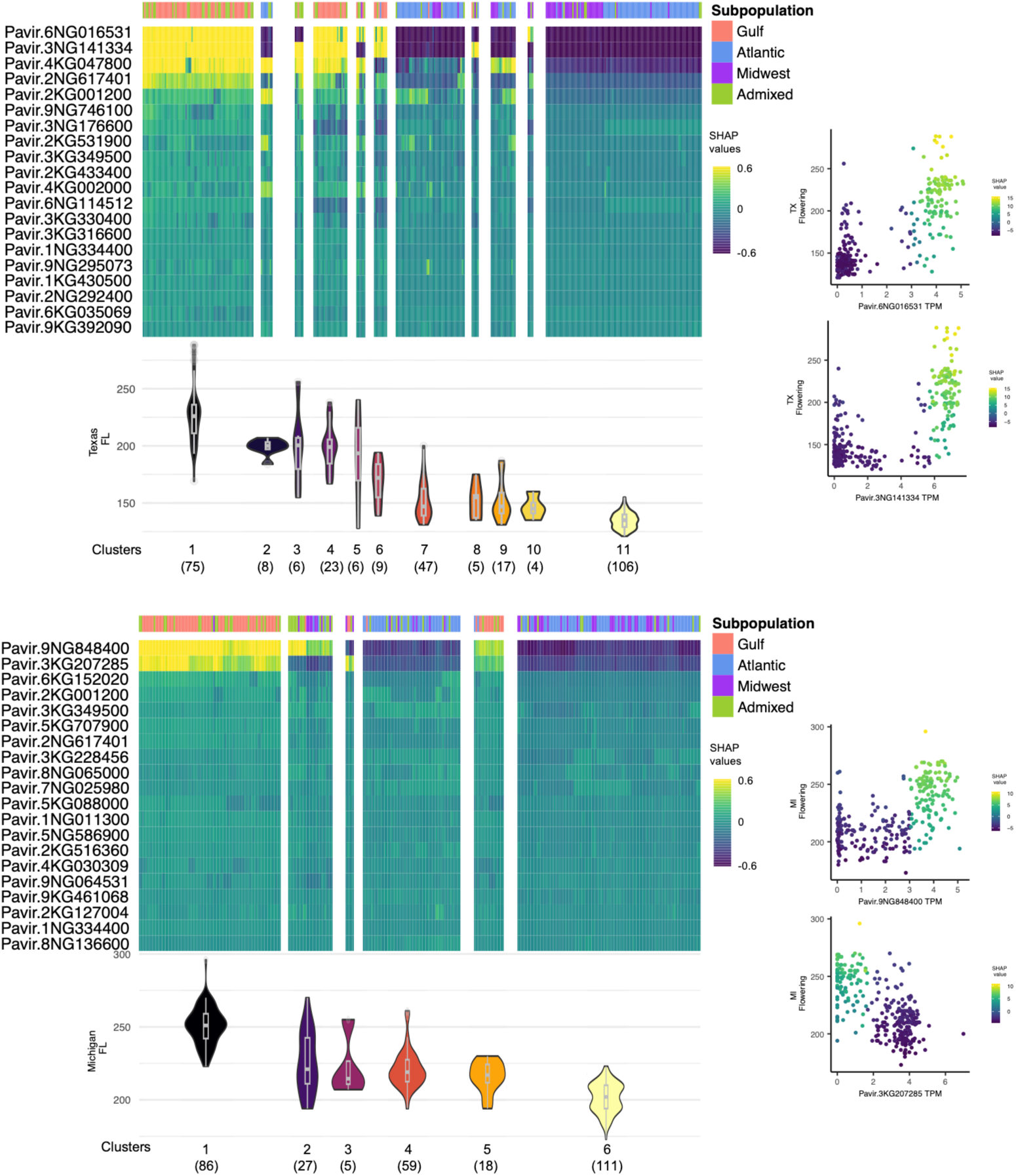
Genotype-dependent effects of top transcriptomic predictors of flowering time in TX (top), MI (middle), and difference models (bottom).

**Supplementary Figure 5.**
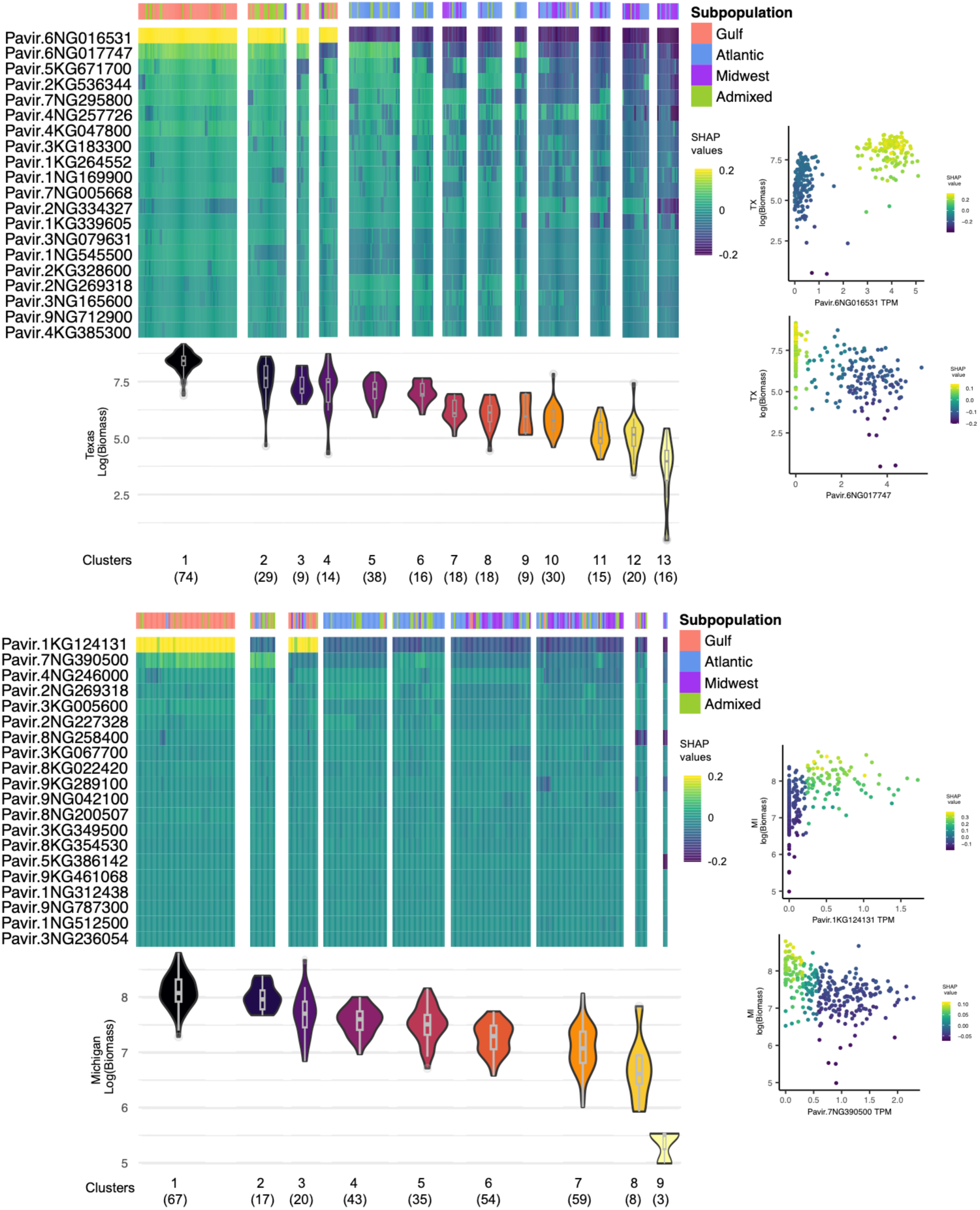
Genotype-dependent effects of top transcriptomic predictors of biomass in TX (top), MI (middle), and difference models (bottom).

**Supplementary Figure 6.**
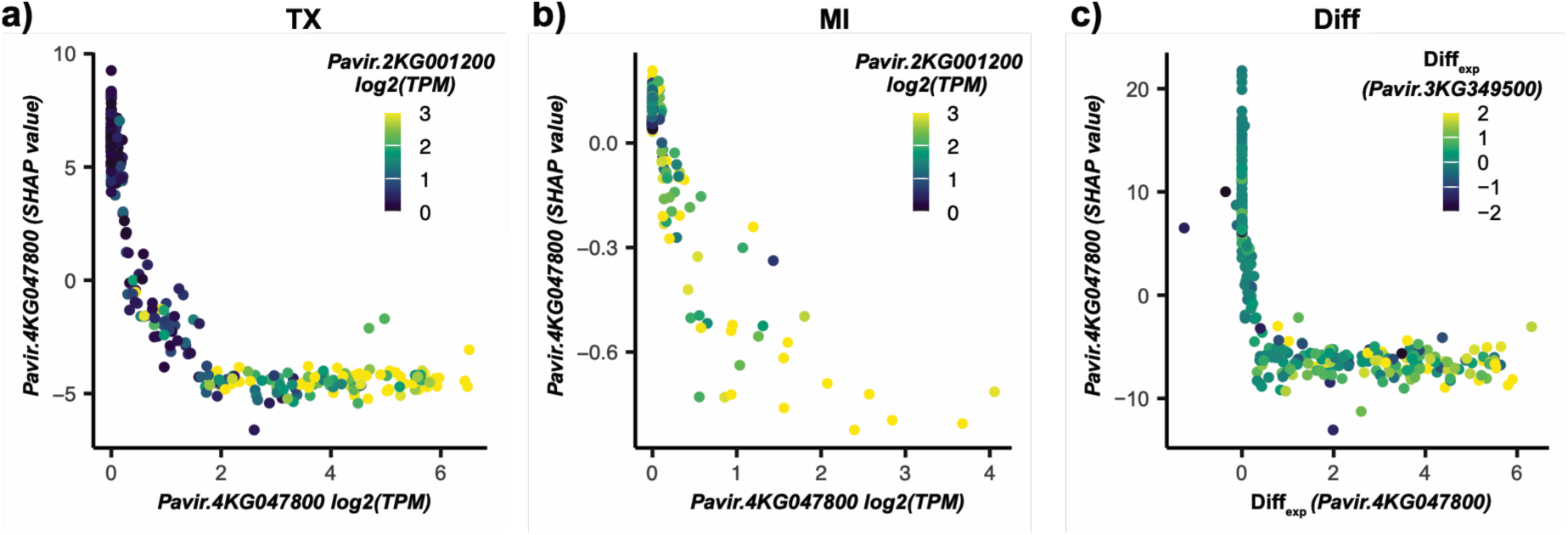
Gene–gene interaction profiles for benchmark flowering time genes, including FT and MADS-box genes, in TX (a), MI (b), and two FT genes in the difference model (c).

## Supplementary tables

***Supplementary Table 1.*** Model training/testing performance and feature selection summary across datasets.

***Supplementary Table 2.*** Feature importance scores and benchmark gene annotations

***Supplementary Table 3.*** Switchgrass homologs of benchmark genes in Arabidopsis and rice for flowering time and biomass

***Supplementary Table 4.*** Candidate biomass genes identified through QTL and GWAS in switchgrass

***Supplementary Table 5.*** Top 100 interacting features identified through SHAP interaction values for flowering time

***Supplementary Table 6.*** Top 100 interacting features identified through SHAP interaction values for biomass

## Notes

### Competing Interest Statement

The authors have declared no competing interest.

### Summary of Updates

The manuscript was revised to improve clarity, align interpretations of the evidence, and address reviewer concerns regarding experimental design, model interpretation, and reproducibility. The title and the abstract were revised to better reflect the study's emphasis on interpretable machine learning, multi-omic integration, and phenotypic plasticity rather than artificial intelligence. Several sections of the Results were rewritten to distinguish analyses based on SNPs, transcriptomic data, and combined models. Statements regarding predictive performance and data complementarity were moderated to avoid over-interpretation, and descriptions of plasticity models were revised to more accurately reflect the observed results. Claims that transcript-difference models outperformed SNP models for flowering time and tiller count were corrected. The Methods and Results sections were expanded to provide a clearer rationale for feature selection and SHAP-based interpretation. Additional text explains how feature selection improved model performance and interpretability, why only a subset of features was used for SHAP interaction analyses, and why SHAP-derived features should be interpreted as predictive associations rather than evidence of causal biological mechanisms. The manuscript now explicitly discusses the effects of linkage disequilibrium, co-expression, and correlated predictors on feature attribution. Revisions were also made to clarify the experimental design. The description of field layout was rewritten, imprecise language was removed, and the honeycomb common-garden design was described in greater detail. The manuscript now explicitly states that only one plant per genotype was evaluated at each site, explains why spatial correction was not feasible, and discusses evidence suggesting limited spatial heterogeneity. Limitations associated with the lack of biological replication for RNA-seq were acknowledged more explicitly. The RNA-seq and transcriptomic analyses were revised to better justify sampling at the pre-flowering stage and to clarify that TX−MI expression differences were used as measures of transcriptomic plasticity rather than for differential-expression testing. Additional discussion was added concerning the compositional nature of TPM-normalized data and the interpretation of transcript-based predictors. Figures were updated with higher-resolution, and legends was revised to clarify correlation analyses, genomic heritability comparisons, SHAP clustering procedures, and transcript-based gene-gene interaction analyses. Additional details were added for statistical models, LD pruning, leaf sampling, training-test stratification, and genomic heritability estimation. Finally, data and code availability were addressed through deposition of TPM expression matrices in Zenodo, addition of Bayesian regression code to the repository, and clarification of RNA-seq processing resources. Several minor corrections, including terminology standardization, FDR adjustment reporting, and correction of the MAF threshold, were also made throughout the manuscript.

